# Ranking protein-peptide binding affinities with protein language models

**DOI:** 10.1101/2024.11.14.623613

**Authors:** Charles Charbel Chalas, Michael P. Dunne

**Affiliations:** Protera Bioscience, Paris, France

## Abstract

In this study we explore the use of protein language models for ranking protein-peptide interaction strength, extending the concept of binary protein interaction classification. We introduce a method that measures and ranks protein binding affinities in an unsupervised manner, eliminating the need for extensive labeled data, structural information, or complex biochemical features. We demonstrate the utility of our approach across five distinct protein-peptide datasets by comparing predicted interaction strength rankings with experimentally derived inhibitory concentration (IC50) values. Furthermore, we discuss limitations encountered during our study and present preliminary findings on extending our approach to more general protein-protein interactions. Finally, we highlight the need for comprehensive datasets specifically designed for ranking protein-protein interactions.

## 1 Introduction

The identification of optimal binding partners for protein-protein and protein-peptide interactions (PPIs) is essential in many bioengineering and therapeutic applications. Computational methods for understanding, identifying, and optimising PPIs have been approached from a variety of angles, each tailored to specific challenges. For example, AlphaFold 3 [2] has shown impressive results in joint protein structure prediction, and RFDiffusion [20] can design binding partners *de novo*. For pairs whose joint structure is known, RosettaDesign [9] and FoldX [19] can be used to optimise interaction interfaces. More traditional methods such as docking, molecular dynamics, and free energy analyses can also be employed using tools such Rosetta [7].

For PPI search, higher-throughput approaches must be considered. Here, large data sets of experimentally determined interactions are often utilised, which typically categorise pairs as either interacting or non-interacting[13][3], for example using yeast two-hybrid screening [3] as evidence. The notion of “interacting” can be quite loose, encompassing everything from short-lived contacts (such as those during signalling cascades), to strong and stable protein complexes [11]. Owing to this, binary classifiers trained on these data sets often have a high margin of error, and at best can be useful for longlisting sets of PPI candidates for a given target protein.

The practical application of PPI studies demands a more nuanced approach, where the strength and likelihood of each interaction is taken into account. Such approaches may be used to rank lists of potential variants of an interacting partner, or to further refine longlists of PPI candidates returned by binary PPI classification searches. A major challenge here lies in the lack of comprehensive databases that quantify interaction strengths: most existing datasets do not offer the detailed interaction strength data necessary for training dedicated machine learning models. This limitation has driven our focus towards using unsupervised learning models, specifically protein language models (PLMs).

In this work, we address the task of ranking potential peptide binding partners for a given protein of interest according to their interaction strengths. Our method builds on work introduced by DiffPalm [10], and utilises ESM2 [6] to rank sets of binding partner variants by considering the relative impacts of their mutation sites on binding affinity. Our method can be applied both to ranking sets of variants suggested by other methods, and for narrowing the search space of semi-rational mutation campaigns by rapidly screening large numbers of potential mutation sites. We demonstrate the utility of the method by comparing its results with five experimental IC50 data sets.

## 2 Methods

### 2.1 Inspiration & motivation

Our method aligns with the framework established by DiffPalm [10], which demonstrated that the reconstruction loss from masked language models (MLMs) can be used to classify protein pairs as interacting or non-interacting. The DiffPalm paper mainly focusses on the MSA Transformer [16], but acknowledges ESM2 [6] as an alternative. To adapt the classification method to ranking scenarios, we focussed on the latter. This choice was driven by the need for a simple and fast solution that bypasses the computationally intensive process of calculating multiple sequence alignments (MSAs). MSAs also are not always available at good depth, especially for shorter sequences such as peptides. ESM2 streamlines the input requirements by directly handling sequence concatenation and focussing on minimizing the masked language modeling (MLM) loss efficiently in evaluation mode. This approach is particularly advantageous in scenarios where rapid and efficient processing is essential.

### 2.2 The MLM loss function

The loss function used in MLMs is designed to assess a model’s ability to correctly reconstruct masked residues during training, through which it learns to understand and interpret complex protein sequences. The MLM loss is defined as follows:

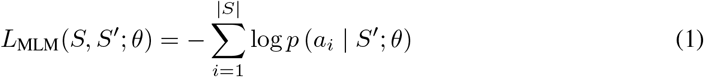

where *S* represents the original sequence, *S′* is the sequence that is inputted to to the ESM2 model, and *a*_*i*_ refers to the actual amino acid at position *i* in the sequence *S*. The term *p*(*a*_*i*_ |*S′*; *θ*) denotes the predicted likelihood of *a*_*i*_ given the input sequence *S′* and model parameters *θ*. In a training scenario, *S′* is equal to *S* but with one or more positions replaced by the `<mask>` token, and *θ* is the set of parameters that is being optimised. We will use ESM2 (version t33-650M-UR50D) in evaluation mode, and so the *θ* term is fixed. We shall therefore refer to the loss between two sequences as simply *L*_MLM_(*S, S′*).

### 2.3 Modifying the MLM loss for scoring position sets

The DiffPalm paper showed that the MLM loss of concatenated and randomly masked MSAs is lower for “correctly” paired (known to interact) sequences than for incorrectly paired ones. For our application we assume that we have a target protein and a binding partner that interacts with it, the aim being to *rank variants* of the partner protein sequence.

To this end, we defined a score that, given a target protein and a chosen reference binding partner, scores a variant by considering only the *positions* for which it differs from the reference. In effect, our score evaluates sets of position in the reference by considering how important they are to ESM2’s ability to reconstruct the paired sequences. For a sequence *S* and a set of indices *I* ⊂ {1, …, | *S*|}, we define the *I*-masked reconstruction loss as:

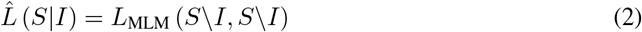

Here *S*\ *I* is the sequence *S* with the amino acids at *I* masked. We define log *p* (`<mask>`| *S*\ *I*; *θ*) = 0, that is, we are calculating the loss on all but the masked positions of the sequence.

### 2.4 Ranking algorithm

To simplify the analysis we restrict the scope of this work to protein-peptide interactions. The more general case of protein-protein interactions is discussed in the Appendices. Our experimental protocol, outlined in Algorithms 1 and 2, takes as input a protein of interest *Q* and a set of peptides {*P*_*i*_ : *i* = 1, …, *N*}. It evaluates the binding potential of each peptide by computing the MLM loss for the concatenation of the protein of interest with each peptide, relative to a reference peptide.

The protocol begins by choosing the reference peptide *P*_ref_ from {*P*_*i*_ : *i* = 1, …, *N*}. The peptide demonstrating the minimal unmasked MLM loss is selected as the reference peptide, serving as an initial hypothesis for the highest affinity interaction (Algorithm 1).

#### Algorithm 1

Reference peptide selection

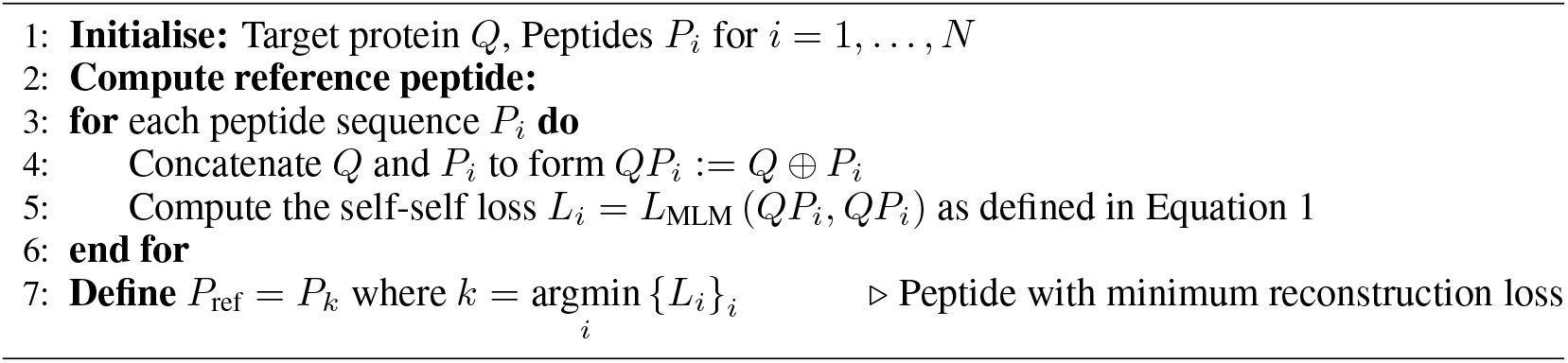

Next, we undertake a differential analysis where each candidate is compared against the reference (Algorithm 2). This comparison involves masking the positions where differences occur between *P*_ref_ and each *P*_*i*_, concatenating them with the target protein *Q*, and reconstructing the result with ESM2. The peptides are then ranked according to their masked reconstruction loss, as defined in Equation 2.

#### Algorithm 2

Ranking position sets

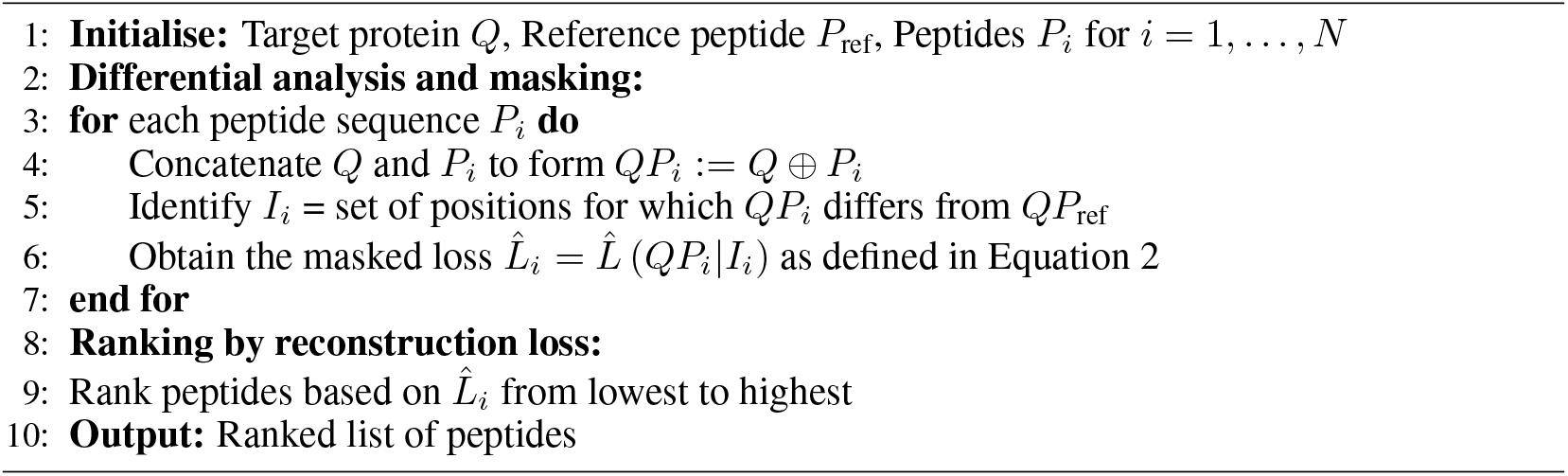

By masking sites, we force the PLM to reconstruct parts of the protein and paired masked peptide. The MLM loss for these masked versions of the peptides provides an indication of the amount of mutual information between *Q* and the *P*_ref_ contained in the masked positions, which can be considered as an indicator of binding affinity. It’s worth noting here that the algorithm does not distinguish peptides with different sequences but which differ from the reference in the same positions (see Discussion). In practice, rather than directly ranking peptides, the approach helps prioritise positions of mutations that are more likely significantly improve binding efficacy.

## 3 Results

We evaluated the performance of our ranking method by comparing its predictions with experimentally derived inhibitory concentration (IC50) values. IC50 measures the effectiveness of a substance in inhibiting a biological function by half and serves as a foundational metric for assessing binding affinity of potential therapeutics against target proteins. We considered five distinct datasets that showcase the method’s potential across different human molecular interactions:

- **HtrA1 and HTRA3**: Serine proteases with a high-temperature requirement, crucial in physiological functions and neurodegenerative diseases like Alzheimer’s disease and CARASIL [4]. We assessed peptides interacting with HTRA1 and HTRA3, with their IC50 values reported in [18] and summarised in Tables A2 and A3. Additionally, we compared our results against the supervised method PPI-Affinity, briefly described in A.4.
- **MDM2 and MDMX**: Both are key regulators of p53 involved in many cancers [15, 12]. Inhibiting their interaction to reactivate p53 represents a therapeutic strategy. Effective 12-residue peptide inhibitors for these proteins are identified in [15] and listed in Table A4.
- **EPI-X4**: An albumin fragment and CXCR4 receptor antagonist with potential in treating HIV, inflammation, and cancer [14][8]. Affinity values for EPI-X4 and 50 derivatives, measured in IC50 nanomolar (nM), were obtained via an antibody competition assay [8]. A comparison with the PPI-Affinity method is also presented to benchmark our results.

We assessed the rankings by visual inspection and also using three ranking performance metrics: normalised discounted cumulative gain (NDCG), precision@K, and Kendall’s *τ*, which are defined in Appendix A.6) and reported for all experiments, with statistics calculated against a random baseline, in Appendix A5.

### 3.1 Case studies: HTRA1 and HTRA3

Our model’s effectiveness is showcased in Figure 1, where it aligns closely with experimental rankings. For HTRA1, our method perfectly matches the top three IC50 rankings, beating PPI-Affinity’s Precision@3 of 0.66. In this case our method’s NDCG of 0.78, while still much better than the baseline, is slightly lower than PPI-Affinity’s score of 0.97. For HTRA3, our model not only scores a perfect Precision@3 but also an impressive NDCG of 0.99, demonstrating robust performance in ranking effectiveness compared to the PPI-Affinity’s NDCG of 0.51, failing notably to rank ‘FGRAV’ as the worst candidate despite its very high IC50 (Table A3).

**Figure 1:**
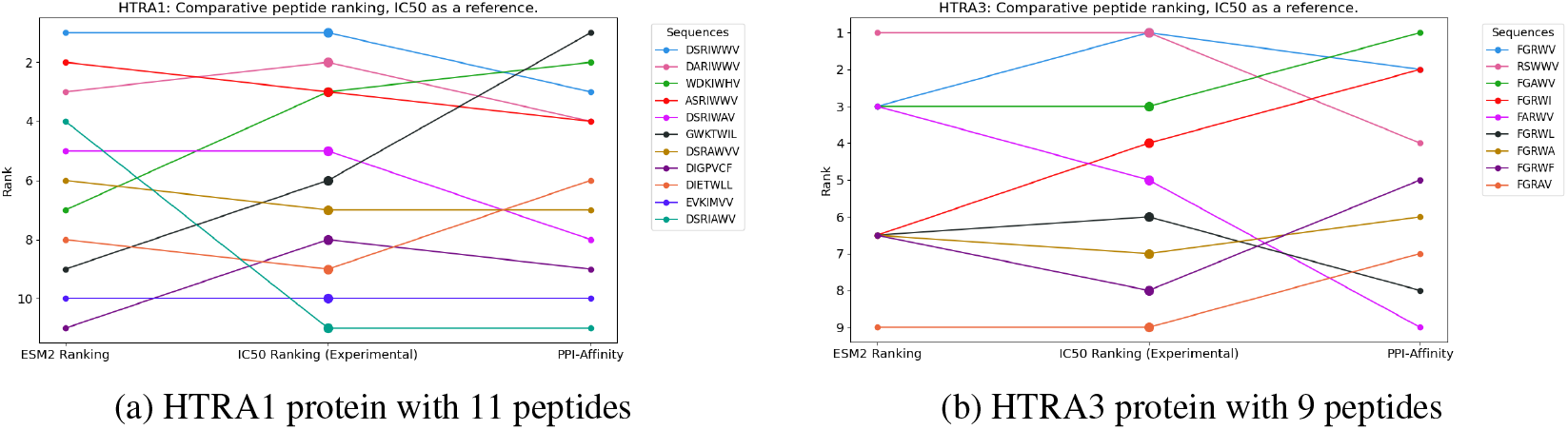
Comparative analysis of HTRAs binding affinities across different methods

### 3.2 Case studies: MDM2 and MDMX

Figure 2 compares IC50 rankings with our predictions for MDM2 and MDMX. For MDM2 (Figure 2a), the ranking was better than baseline (NDCG=0.67) and identified one of the top three peptides, yet the overall results were not satisfying. The results for MDMX (Figure 2b) were more favorable; our method accurately predicted five of the top six peptides, missing only the top one, yielding a Precision@3 of 0.66 and an NDCG of 0.64, highlighting NDCG sensitivity to misrankings top candidates. The incorrect ranking of the inactive peptide 4T6W in both cases highlights the model’s limitations, and we are investigating the resons for this.

**Figure 2:**
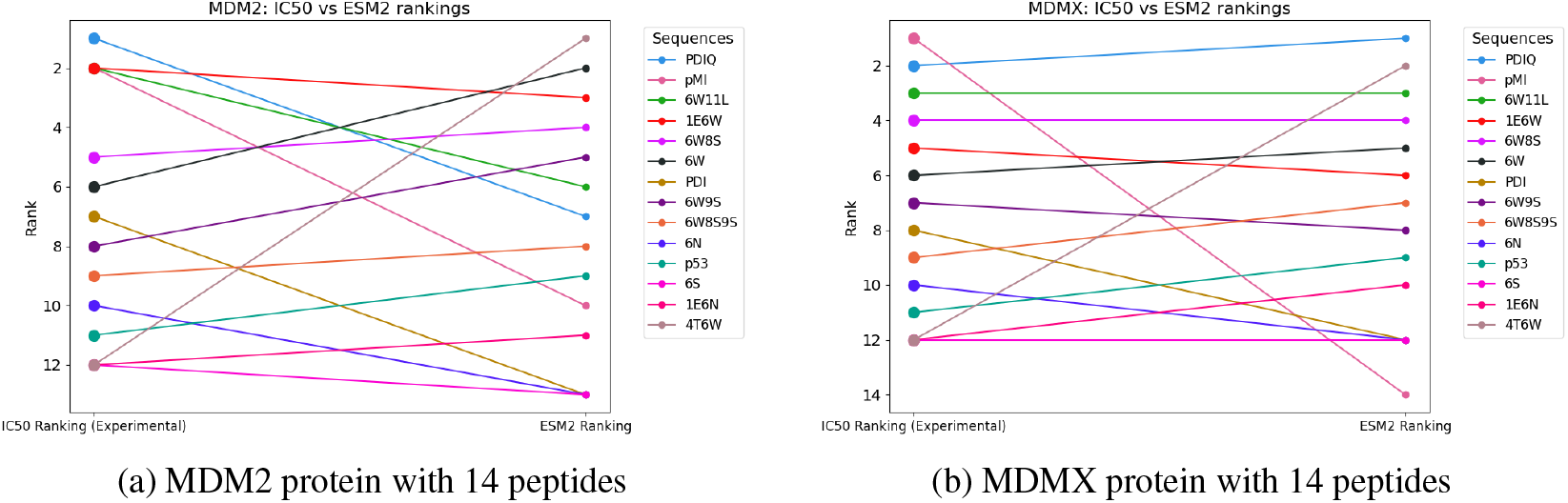
Comparative IC50 vs ESM2 rankings on MDM2 and MDMX proteins

### 3.3 Case study: EPI-X4

For EPI-X4, we grouped the 50 derivatives by peptide length for analysis, as shown in Figure A7. Our method consistently surpassed PPI-Affinity, correctly identifying top candidates for lengths 10 and 11 at position 1, and for length 9 at position 2 (Figures A7b, A7c, A7d). PPI-Affinity, in contrast, ranked these at the bottom. While both methods struggled with the first group of 19 length-12 peptides (Figure A7a), our method achieved an overall NDCG of 0.9 compared to PPI-Affinity’s 0.79. Interestingly, PPI-Affinity’s performance was below that of a random ranker in terms of NDCG, despite a higher Precision@3 of 0.87.

## 4 Discussion

### 4.1 ESM2 can be used to rank protein-peptide binding affinities

Our study indicates the effectivenes of PLMs in accurately predicting relative interaction strengths, aligning well with experimental IC50 values. This method overcomes the need for labelled datasets and complex feature engineering, providing a streamlined approach for examining protein interactions. Our method is fast and can be used to rank and filter large sets of possible peptide mutants to reduce the number of candidates sent to the lab.

Our method ranks sets of positions relative to a chosen reference peptide. The obvious drawback here is that it cannot distinguish mutations that occur at the same positions. More work needs to be done to extend the method to these scenarios. Knowing how certain positions affect binding affinity is however useful on its own: it allows us to massively reduce the search space of potential mutations by indicating only those positions which are likely to yield successful mutants. This information can be leveraged in mutation campaigns to produce improved peptide candidate sets.

### 4.2 Our method showed some limitations

In evaluating MDM2 and MDMX, the ESM2 model showed limitations, particularly with inactive peptides. It correctly ranked two out of three inactive peptides as low, but significantly misranked 4T6W. This inconsistency highlights the model’s occasional difficulty in identifying non-active peptides, suggesting that further investigation is needed to determine if a solution is possible.

### 4.3 PPIs and docking show promising preliminary results

While our study primarily focused on quantifying protein-peptide binding affinities, we expanded our analysis to protein-protein interactions and docking scenarios, achieving promising results as detailed in Appendices A.2 and A.3. However, the lack of reliable protein-protein interaction ranking datasets is a major challenge. We encourage the community to develop these datasets, as they are crucial for accurately predicting and validating effective interaction candidates in practical applications.

## 5 Conclusion

In this study, we validated the use of unsupervised protein language models (PLMs) for ranking protein-peptide pairs according to their binding affinities, with promising extensions to protein-protein interactions. Our approach was tested across various datasets, effectively ranking interaction strengths, as evidenced by comparisons with experimental IC50 values and established computational tools. These findings illustrate the potential of PLMs for assessing protein interactions without extensive supervised training or resource-intensive computational biology methods. Our analyses underscore the need for specialised PPI datasets tailored for ranking to enhance their applicability in real-world scenarios, setting a foundation for future advancements in comparative protein interaction analysis.

## A Appendix / Supplemental material

### A.1 Code availability

Our code is available at https://github.com/proterabio/ranking-protein-peptides

### A.2 Ranking protein-protein datasets

While the main focus of this report has been protein-peptide interactions, the initial scope of our study was to rank binding affinities for a protein of interest with any group of potential protein candidates. In this section, we aim to expand on the work above by testing our method on protein-protein affinities.

Human DPP4 (Dipeptidyl Peptidase-4) is a known receptor for the Middle East Respiratory Syndrome Coronavirus (MERS-CoV) RBD protein, facilitating the virus’s entry into host cells. To investigate the impact of specific DPP4 mutations on its binding affinity to RBD, we utilised a data set previously analysed using the Mutabind2 computational tool (a supervised method requiring structure as input introduced in [22]). This analysis, which was detailed in a prior study [1], employed custom calculation parameters based on the Gibson free energy (ΔΔ*G*) equation, leading to varied predictions on the effect of mutations on protein-protein interactions (PPIs), as can be seen in Table A1.

**Table A1:** DPP4 – MERS-CoV Binding Affinity calculated by Mutabind2

| Mutation | $\Delta\Delta G$ (kcal/mol) | Predictive Mutation Effect |
| --- | --- | --- |
| N229I | 0.47 | Destabilizing |
| K267N | 1.9 | Destabilizing |
| K267E | 3.63 | Destabilizing |
| T288P | 0.03 | Destabilizing |
| L294V | 2.0 | Destabilizing |
| L295L | 0.94 | Destabilizing |

Similar to the analysis conducted in Section 3.1, we compared the rankings produced by the Mutabind2 tool with those derived from our ESM2-based method. Although the rankings from these two tools do not align perfectly, there is notable agreement in key areas (Figure A3). Specifically, both our method and Mutabind2 identify the same Top1 and Top2 candidates, highlighting a consistent recognition of these interactions as the most significant. Additionally, both approaches concur on ranking the K267E mutation as the least effective candidate, further validating the reliability of our predictive model against established computational tools.

**Figure A3:**
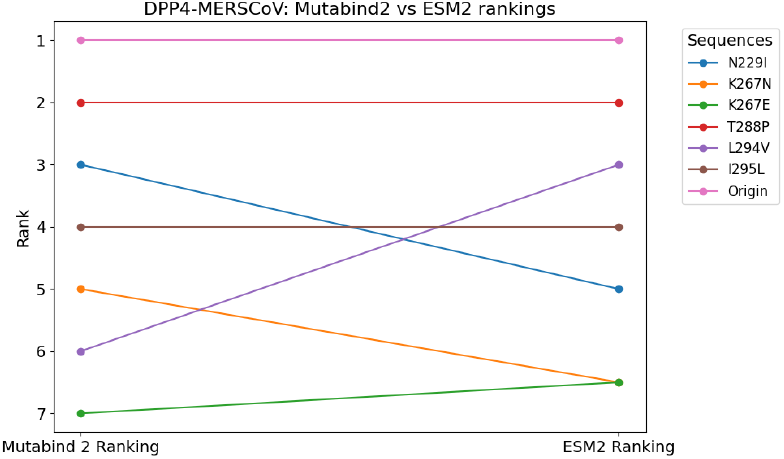
Mutabind2 vs ESM2 ranking on MERS-cov proteins with several DPP4 variants

**Figure A4:**
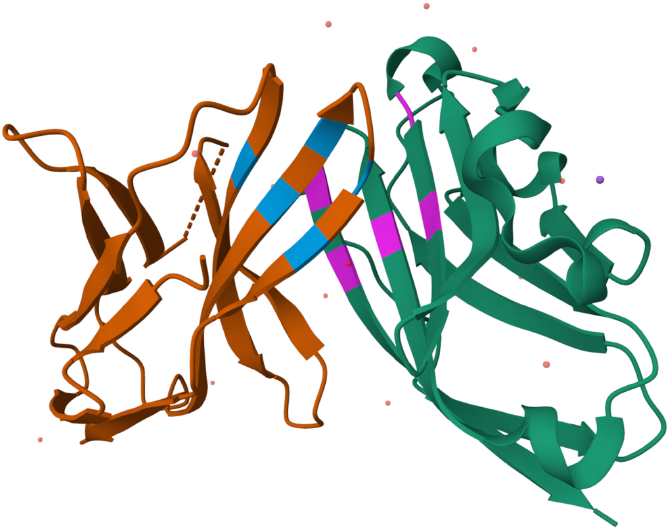
Interaction zone between PD1 and PDL1: The blue region of PD1 engages with the pink region of PDL1 as identified in [21]

### A.3 Case study: PD1-PDL1 in a docking scenario

Though it was not the main focus of this study, we were intrigued to explore whether our approach could be extended to docking applications. The interaction between programmed cell death protein 1 (PD1) and its ligand PDL1 is a critical regulatory mechanism in the immune system, playing a pivotal role in the modulation of immune responses and the maintenance of self-tolerance. [21] studied this interaction and identified the amino acids that actually interact, as can be seen in Figure A4. We aimed to use ESM2 to investigate the potential binding site of PDL1, assuming that the binding site of PD1 was unknown. Instead of calculating the overall reconstruction loss for both proteins after masking patch residues on PDL1, as we had done in previous experiments, we focused solely on the reconstruction loss of those masked residues in this docking scenario.

We recognise that the task of docking involves complexities beyond simple interaction prediction, notably the necessity to consider solvent-accessible amino acids. The presence of amino acids that can theoretically interact does not guarantee their interaction in a physical context: both must be accessible on the protein’s surface for an interaction to be feasible. In this experiment, we identified exposed residues using SASA calculations obtained using BioPython [5]. We proposed N=44 sets of k=5 positions on PDL1 by identifying surface amino acids from its PDB file and then choosing combinations of amino acids that are all within a distance threshold (8Å) from each other. For each identified position set, we calculated the reconstruction loss to assess the likelihood of interaction between PD1 and PDL1 at these specific positions. A lower loss suggests a greater probability of interaction, helping to pinpoint potential critical interaction zones between the two proteins.

Figures A5 and A6 demonstrate that the amino acids identified by our method in both the Top1 and Top2 candidates are very near the experimentally determined ground truth regions, affirming the meaningfulness and accuracy of our predictions. The experimentally validated ground truth was ranked a close third, following these top two candidates. Our method is quick and straightforward in cases where a PDB is available, which may be obtained via structure prediction. Further exploration is necessary to determine if we can proceed using sequence alone.

**Figure A5:**
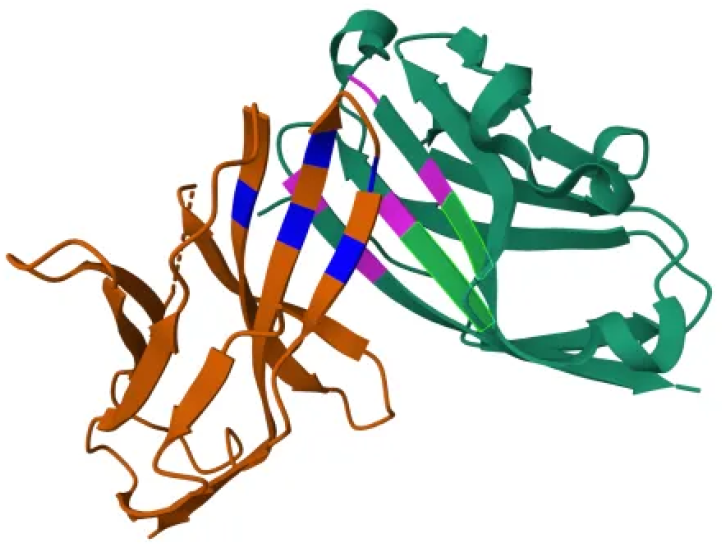
Top-Ranked candidate highlighted in green according to our method. Blue and pink amino acids represent correct interactions according to [21]

**Figure A6:**
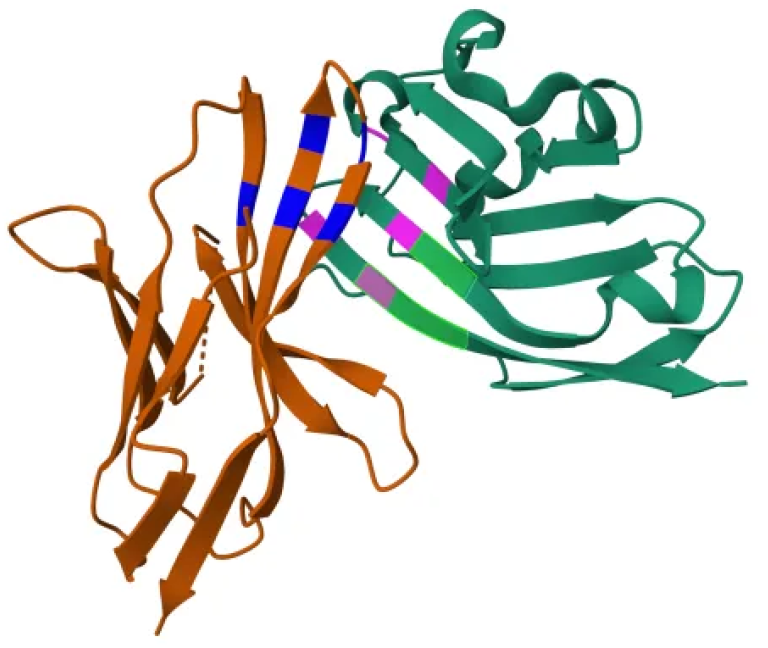
Second-Best candidate highlighted in green according to our method. Blue and pink amino acids represent correct interactions according to [21]

### A.4 Binding Affinities of Peptides to the HTRAs and a summary for PPI-Affinity method

Tables A2 and A3 summarise the IC50 values for HTRA1 and HTRA3, as reported by [18] and as discussed above in Section 3.1.

For these proteins, we compared our results against the PPI-Affinity method, a recent supervised approach detailed in [17]. This method is currently available only through a web server. Unfortunately, we were unable to access the server for live comparisons, so our comparative analysis uses results directly copied from their publication. This method utilises a complex feature set consisting of 23,040 descriptors for each of the 1,149 protein-peptide complexes it analyzes, and employs a Support Vector

Machine (SVM) for training. By comparing these published results, we can benchmark the efficacy of our unsupervised approach against established experimental data and a sophisticated supervised learning model.

**Table A2:**
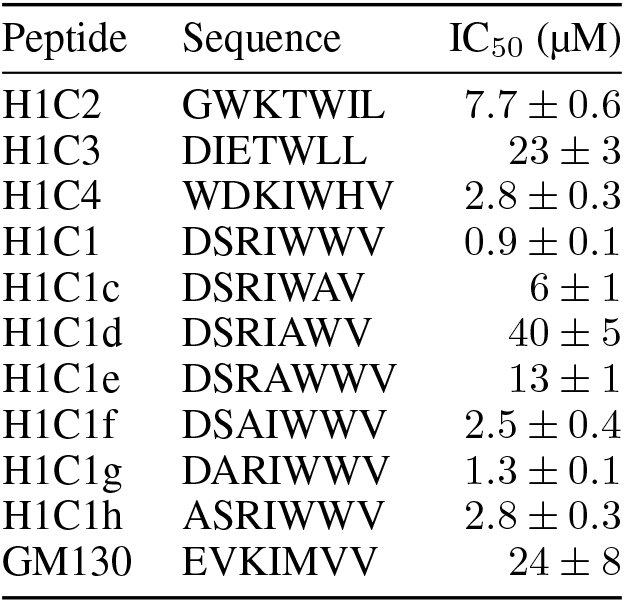
IC_50_ values for synthetic peptides binding to HTRA1-PDZ as reported in [18]

**Table A3:** IC_50_ values for synthetic peptides binding to HTRA3-PDZ as reported in [18]

| Peptide | Sequence | IC<sub>50</sub> (μM) |
| --- | --- | --- |
| H3C2 | RSWWV | 0.6 ± 0.1 |
| H3C1 | FGRWV | 0.6 ± 0.1 |
| H3C1a | FGRWI | 1.0 ± 0.1 |
| H3C1b | FGRWL | 2.9 ± 0.3 |
| H3C1c | FGRWF | 7.7 ± 0.8 |
| H3C1e | FGRWA | 3.5 ± 0.3 |
| H3C1f | FGRAV | 270 ± 110 |
| H3C1g | FGAWV | 0.9 ± 0.1 |
| H3C1h | FARWV | 1.1 ± 0.2 |

### A.5 Binding Affinities of Peptides to MDM2 and MDMX

Binding Affinities of Peptides to MDM2 and MDMX are presented in Table A4. IC50 values are reported in [15].

**Table A4:**
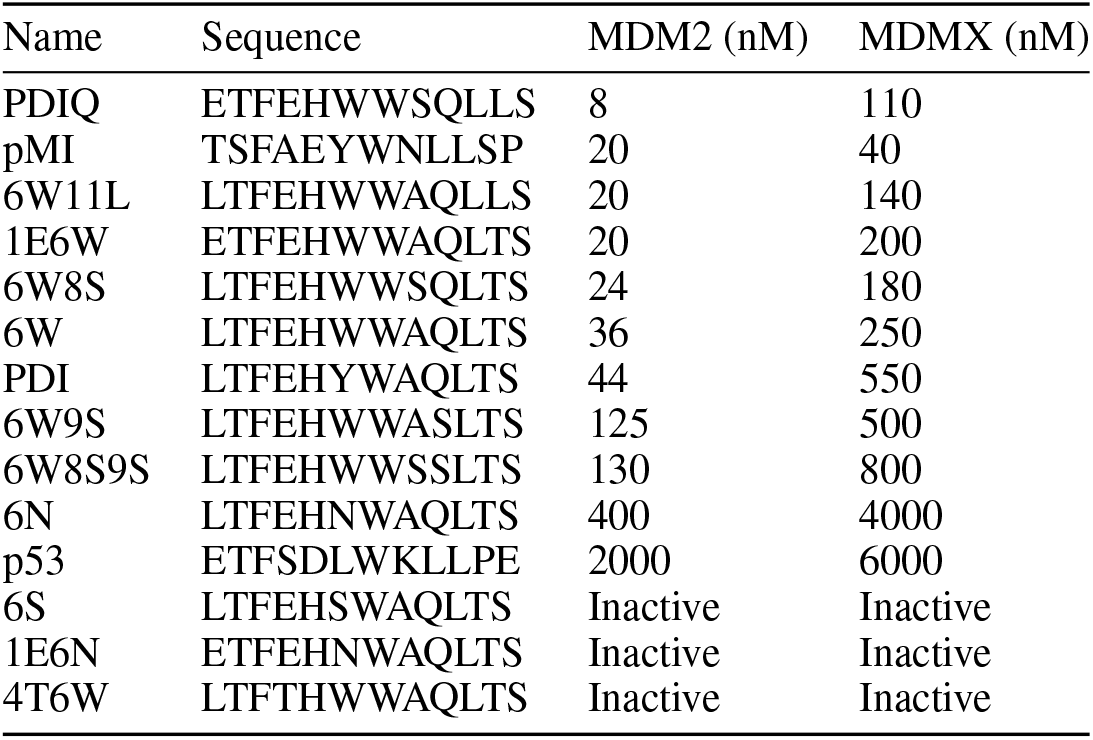
Binding Affinities of Peptides to MDM2 and MDMX. Values from [15]

### A.6 Metrics and baseline evaluation

#### Normalized Discounted Cumulative Gain (NDCG)

is a measure of ranking quality that evaluates the order of a list of predictions based on their relevance. It gives higher importance to hits at top ranks and discounts the value of hits at lower ranks. NDCG is particularly useful in scenarios where the highest ranked results are more significant than those lower down the list. It is calculated using the formula:

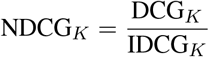

where DCG_*K*_ (Discounted Cumulative Gain) is:

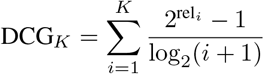

**Table A5:**
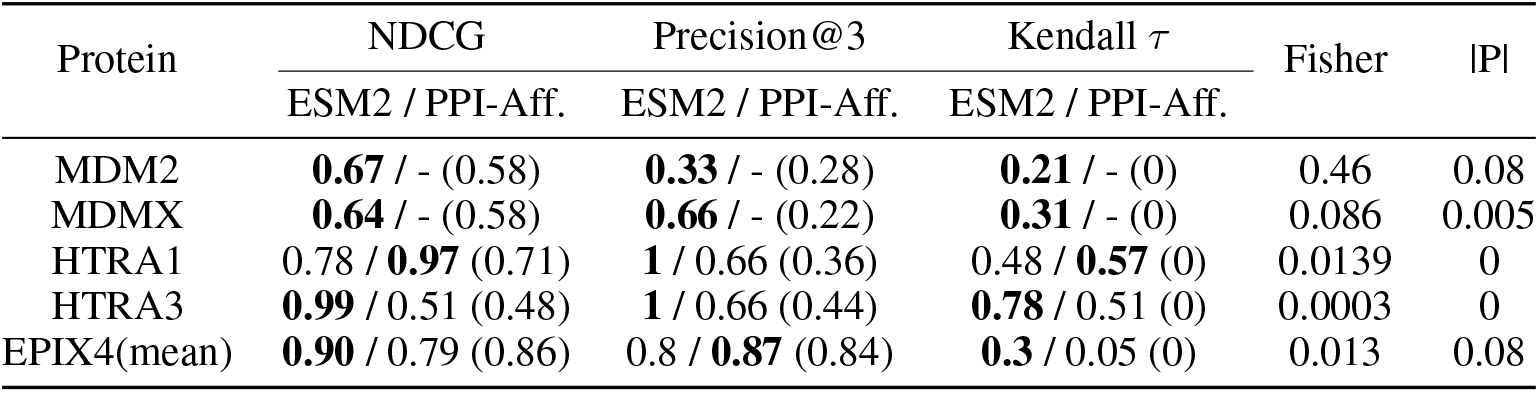
Comparison of NDCG, Precision@3, and Kendall *τ* for ESM2 rank and PPI-Affinity. For our ESM2-based method, we also report aggregated p-values using the Fisher method and |P| score, defined in Appendix A.6. Baseline values are in brackets, with *τ* = 0 the theoretical baseline for *τ* and the others calculated by taking the mean across 10000 random orderings. In all cases, the probability |P| of observing equal or better results in all three metrics was extremely low. The fact that the method struggled on MDM2 is apparent in its Fisher combined p-value, and the result for MDMX is also borderline. More investigations need to be performed to understand these cases.

Here rel_*i*_ is the relevance score (in this case the values being measured, for example the ESM2 loss or the PPI-Affinity score) of the item at position i and IDCG_*K*_ is the ideal DCG, the maximum achievable DCG with the same set of relevance scores but in the perfect ranking order.

##### Precision@K

is a metric used to evaluate the accuracy of the top *k* predictions. It measures the proportion of relevant items found in the top *k* recommendations of the ranking model. This metric is crucial when the goal is to identify a small number of highly relevant items from a larger set, making it ideal for assessing the effectiveness of ranking proteins or peptides in bioinformatics. It is defined as:

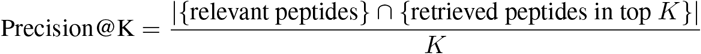

#### Kendall Tau Rank

(*τ*) is a non-parametric statistic used to measure the ordinal association between two variables. It evaluates the number of concordant and discordant pairs to determine the correlation. Values range from −1 (perfect disagreement) to +1 (perfect agreement), with 0 indicating no correlation. This metric is useful for assessing monotonic relationships and is robust against outliers.

Table A5 shows a comparison of our method against PPI-Affinity and a baseline, which is a random ordering run 10,000 times with its scores averaged resulting in Fisher coefficient and |P| value. Fisher’s method, also known as Fisher’s combined probability test, was developed by Ronald Fisher and is used to combine results from several independent tests related to the same overall hypothesis (H0). The |*P*| value, on the other hand, represents the proportion of times the random ranker outperformed our method across all three metrics used in this study.

### A.7 Case study: EPI-X4 affinity to CXCR4 derivatives

**Figure A7:**
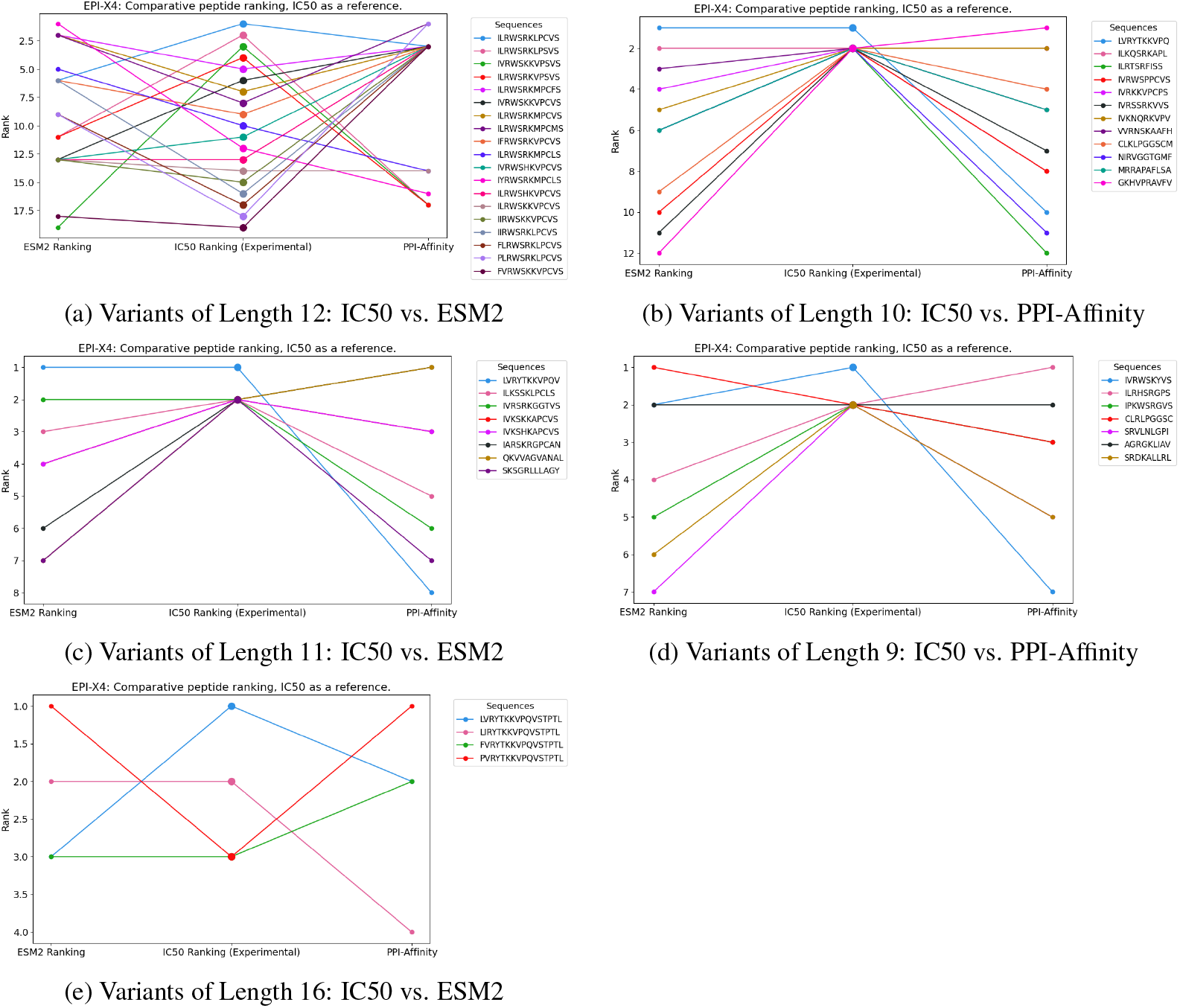
Comparative Analysis of EPIX4 bindings to variants of CXCR4 using our method, PPI-Affinity paper and IC50 values (published in [8])

